# C57BL6 mouse substrains demonstrate differences in susceptibility to the demyelinating effects of Cuprizone toxin

**DOI:** 10.1101/2020.12.17.423334

**Authors:** Elaine O’Loughlin, Binod Jacob, Gonzalo Zeballos, Philip Manfre, Anjali McCullough, Nicholas Thomas Gatto, Takayuki Tsuchiya, Anna-Mari Karkkainen, Kimmo Lehtimäki, Juha Kuosmanen, Thomas W. Rosahl, Geoffrey B. Varty, Matthew E. Kennedy, Christian Mirescu, Sophia Bardehle

## Abstract

Advances in our understanding of cellular functions and phenotypes in the brain rely on technically robust experimental *in vivo* models with face validity towards human disease. The cuprizone toxin-induced demyelination model is widely used to investigate pathophysiological mechanisms of demyelinating and remyelinating phases of multiple sclerosis. The C57BL6 mouse is a common inbred strain used as the genetic background for genetically engineered and congenic mice. Substrains of C57BL6 mice sourced from distinct vendors are often treated as equivalent in research studies. Here, we demonstrated that an alternative dosing approach via oral gavage with a well-tolerated, lower dose of cuprizone resulted in significant differences in C57BL/6NTac (Taconic) over C57BL/6J (Jax) mice. With consistent dosing of cuprizone for 5 weeks, body weights were significantly affected in C57BL/6NTac versus C57BL/6J mice. DT-MRI showed significant demyelination in white matter regions in the C57BL/6NTac mice. Concomitantly, histology analysis illustrated increased microgliosis and proliferation in C57BL/6NTac compared with C57BL/6J mice. These observations suggest that the C57BL/6NTac substrain of C57BL6 mice is more vulnerable to cuprizone challenge. Genetic factors along with breeder source appear to influence susceptibility to cuprizone toxin. Thus, the awareness of the limitations of *in vivo* models in addition to informed decision making on the appropriate background substrain can greatly improve sensitivity and reproducibility of results and use for evaluating investigational therapeutics.

## 1. Introduction

The C57BL/6 mouse strain has become the most frequently used inbred strain in biomedical research with over 44,000 articles in PubMed documenting its use. It has several advantages such as ease of introducing transgenic mutations, breeding, disease modelling and efficacy studies. However, numerous studies have demonstrated physiological differences between mouse strains and even between the same strain bred at different commercial vendors and for this reason, it is very important to be informed on the strain and substrains of choice (Mekada et al., 2009, Bryant, 2011, Fontaine and Davis, 2016). The application of mouse models to the study of neurological disease has proven to be a highly valuable approach to understanding disease pathophysiology and for the evaluation of investigative therapies. Moreover, *in vivo* models allow the study of pharmacokinetics and pharmacodynamics, drug metabolism, efficacy, safety and the development of biomarkers (Bai et al., 2016). Genetic engineering or chemical-toxin treatments are common manipulations used to induce disease relevant phenotypes that allow controlled investigation of specific gene function and therapeutics of interest. Selection of appropriate mouse strains is a key aspect in designing a study that can support sound scientific interpretations. Unappreciated biological differences between different mouse strains, such as differential sensitivity to genetic or chemically induced disease phenotypes could mislead researchers in interpreting gene or pharmacological dependent outcomes. Thus, choosing an appropriate murine strain for any disease model is critical and requires validation of multiple variables. Here, we carried out a deep characterization of the effects of cuprizone (cpz)-induced demyelination in two commonly used substrains of C57BL/6 mice: C57BL/6J (The Jackson Laboratory) and C57BL/6NTac (Taconic Laboratories). To our knowledge, this is the first study to evaluate the susceptibly of substrain differences in two widely used strains of C57BL6 mice in response to cpz-toxin by longitudinal brain imaging and histopathology

Studying innate immune biology in the brain can be impacted by technical variables as illustrated by recent publications, identifying stronger protocol than biology dependence of transcriptomic profiles of microglia (Gosselin et al., 2017, Krasemann et al., 2017, Dubbelaar et al., 2018). Microglia are the brains phagocytes that are largely responsible for the clearance of debris, including myelin debris and play critical roles in demyelinating diseases. MS is characterized by damage of myelin associated with infiltrating immune cells and inflammation that leads to progressive axonal degeneration and synaptic loss, coupled with the recruitment of resident microglia to sites of myelin injury (Lassmann, 2001, O’Loughlin et al., 2018). It is a multifaceted heterogenous disease involving adaptive and innate immune mechanisms, thus there is no single animal model that faithfully represent the entirety of the human disease (Ransohoff, 2012, Lassmann and Bradl, 2017). Experimental autoimmune encephalomyelitis (EAE) is a CD4^+^ T-cell mediated autoimmune disease model and is robustly used to study innate and adaptive immune components and their interactions with the central nervous system (CNS) (Ryu et al., 2018). In contrast to autoimmune models, the cuprizone model is a toxin-induced, reversible demyelination model with selective cell death of myelin-generating oligodendrocytes leading to demyelination associated with a characteristically strong microglia response but minimal blood brain barrier (BBB) damage and peripheral immune cell infiltration (Berghoff et al., 2017). Cessation of cuprizone toxin treatment leads to spontaneous remyelination While this model captures aspects of the MS pathophysiology, it is a broadly utilized pre-clinical *in vivo* tool to study cell biological and molecular pathways underlying de- and remyelination processes. A key advantage of this model is the ability to study the function of innate immune biology effects on the pathophysiology of demyelination/remyelination in the CNS without confounding peripheral infiltrates such as T-cells (Praet et al., 2014, Hillis et al., 2016).

Cuprizone (cpz) damages oligodendrocyte homeostasis by chelation of copper and induction of oligodendrocyte apoptosis with subsequent demyelination (Cammer and Zhang, 1993, Skripuletz et al., 2011). Since oligodendrocyte precursor cells are resistant to cpz toxicity, remyelination commences following cuprizone washout. Demyelination/remyelination is largely observed in white matter rich regions such as the corpus callosum (CC) and is accompanied by gliosis (microgliosis and astrogliosis) and neurodegeneration (Praet et al., 2014). Cpz supports a focus on mechanisms of demyelination-and remyelination, the availability of validated biomarkers and histological analyses to score disease and the rapid onset and resolution phases allows for the study of novel interventions directed at microglia or oligodendrocyte biology for example. By comparing the two most widely used C57BL/6 substrains, we show that 6NTac mice are more vulnerable to cpz-induced demyelination in the CC in comparison to 6J mice over a 5-week dosing paradigm. Murine genetic background greatly influences the susceptibility of mice to cpz-induced damage, and that the influence of mouse strains should be taken into consideration in designing experiments using the cuprizone model. Understanding and communicating mouse sub-strain differences and variability to injury responses are critical to not only the reproducibility of early discovery studies within the scientific community but are also important for effectively choosing efficacy studies for pharmacological development for neurological diseases. Substrain differences of C57BL6 mice were examined as some genes of interest e.g. TREM2, are bred onto different backgrounds to study neurodegeneration for example, with reference database can be found online https://www.alzforum.org/research-models. It is important to choose the appropriate strain background in order to study the biology correctly and for drug target validation in a disease context.

## 2. Results

### 2.1 Differences to cpz response in C57BL/6NTac and C57BL/6 Jmouse strains

Body weight (BW) loss is a characteristic trait of the cpz-induced demyelination model. BW was recorded daily for 35 days until end of in life study and there were no body weight differences recorded at baseline between strains (Figure 1B, C). BW was presented as daily gram (g) weight (Figure 1B) and percentage change (%) from pre-dose BW (Figure 1C). Cpz administration resulted in significant reduction in BW over time between the two strains. Over the course of the 5-weeks, cpz dosing resulted in 6NTac mice have a significant decrease BW compared to 6J mice over varying days (g and % change) (Figure 1 B and C). The percentage BW change from day 0 was taken to eliminate differences from starting weights and give an accurate value of the BW loss. Comparison of strain in cpz dosing only illustrated significantly changes in BW change (%) from baseline between the groups (Figure 1B). The data indicates that C57BL/6NTac mice are more vulnerable to cpz toxin challenge in comparison to C57BL/6J mice.

**Figure 1:**
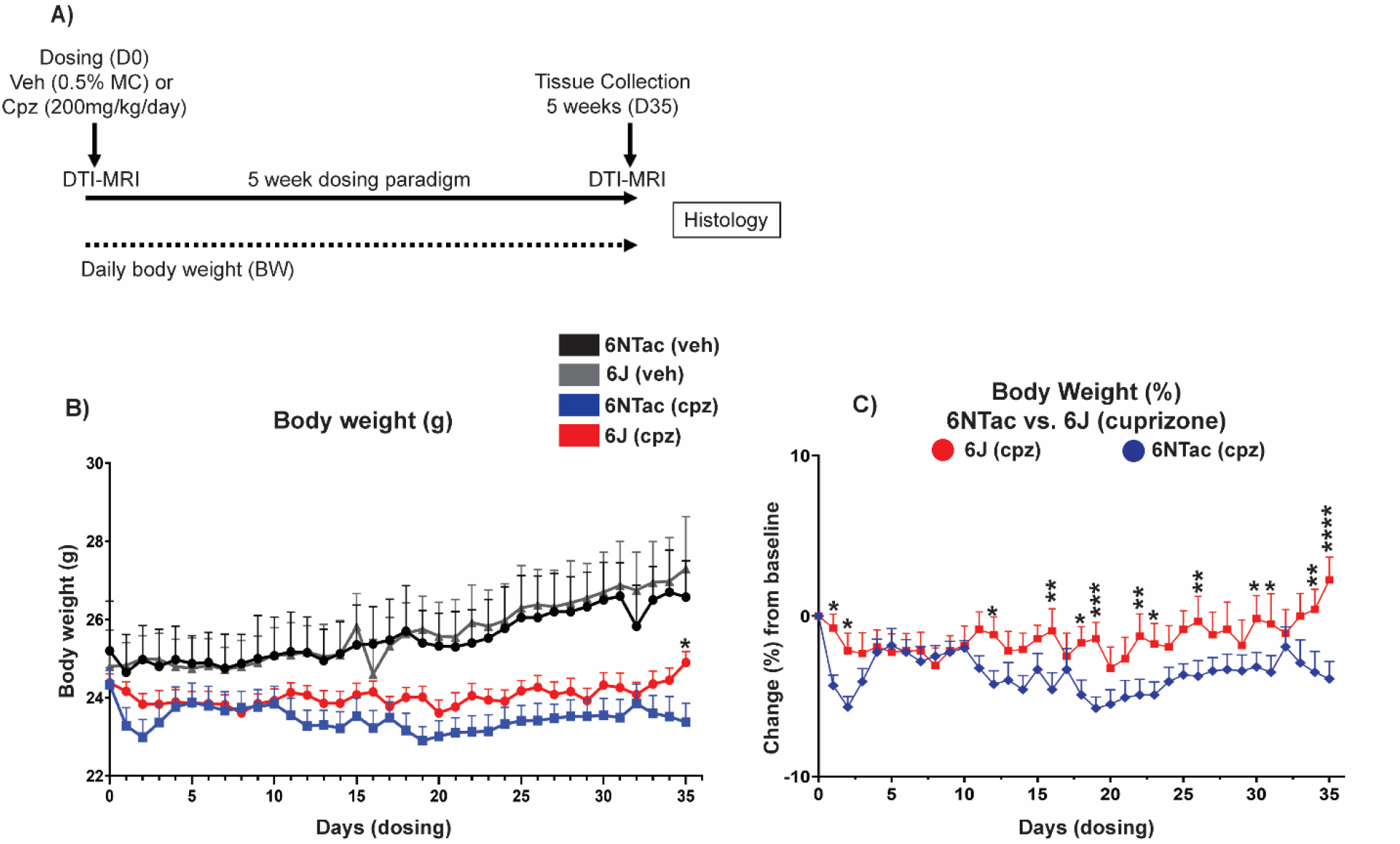
A) Schematic of experiment design. Animals were dosed with Cpz (cpz; 200mg/kg/day) or vehicle (veh) twice daily for 5 weeks (35 days). Body weights were taken daily at consistent time of the day. Diffusion Tensor Imaging (DTI) acquired at 1 day prior to starting dosing (baseline) and on the final day of dosing at D35. B) Mean body weight (g) monitored during cpz dosing from baseline (D0) to Day 35. C) Percentage body weight (%) change from baseline in C57BL/6NTac and C57BL/6J mice after cpz treatment over 5 weeks (D0; 0%). Data represented mean BW ±SEM. Statistical significance (*, p<0.05; **, p<0.01; ***, p<0.001) was evaluated with two-way ANOVA Tukey’s multiple comparisons. Animal groups; n= 12 6J+cpz, n= 12 6NTac+cpz and n=4 6J+vehicle, 6NTac+vehicle per group.

### 2.2 Effect of cpz oral gavage on demyelination by DT-MRI imaging

DT-MRI is sensitive to microstructural tissue properties and can be used in pre-clinical research and clinical work to explore white matter anatomy and architecture structure *in vivo.* DTI was carried out on all mice (cpz and vehicle treated) at baseline and 5 weeks (Figure 2A). We used this *in vivo* imaging method to assess structural abnormalities in CC subregions in all animal groups. Fractional anisotropy (FA) values were extracted to show the extent of demyelination and loss of structure coherence within the white matter regions, specifically subregions of the CC. Labelled subregions of the CC are outlined in Figure 2A. After 5 weeks dosing, C57BL/6NTac and C57BL/6J cpz treated animals exhibited significant differences in FA in measured subregions of the CC, in comparison to their corresponding vehicle controls (Figure 2A-C). Across subregions of the CC studied, including forceps minor (FMI), external capsule (ec), body of CC (Bcc), genu of the CC (Gcc) and the splenium (Scc), FA measurements were significantly reduced in C57BL/6NTac mice in comparison to C57BL/6J mice (Figure 2C). Representative images of FA analysis between C57BL/6NTac and C57BL/6J cpz treatment are seen in Figure 2A&B. The inset panel (Figure 2B) depicts example regions, the genu and body of the CC with the FA image. The MRI image clearly shows a loss of FA signal in this region in C57BL/6NTac in comparison to C57BL/6J and this is quantified (Figure 2B; red arrow & 2C). This suggests that C57BL/6NTac have heightened sensitivity to cpz toxicity resulting in increased demyelination and structural derangement of the CC. Deeper white matter structures such as the internal capsule, optic tract and cerebral peduncle exhibited less damage to cpz challenge. No changes were observed by FA in these regions *(data not shown).* We also extracted the mean diffusion values (orientation independent measure of average diffusion) from generated tensor parameters. No differences were recorded in the mean diffusion between C57BL/6NTac and C57BL/6J animals treated with cpz for the body of the CC, splenium of the CC or the external capsule, respectively *(Supplementary Figure 1).* However, the forceps minor of the CC and the genu of the CC showed increases in mean diffusion in C57BL/6NTac but not C57BL/6J cpz treated animals (between baseline and 5 weeks treatment) further illustrating the greater susceptibility of C57BL/6NTac to cpz challenge in comparison to C57BL/6J *(Supplementary Figure 1*).

**Figure 2:**
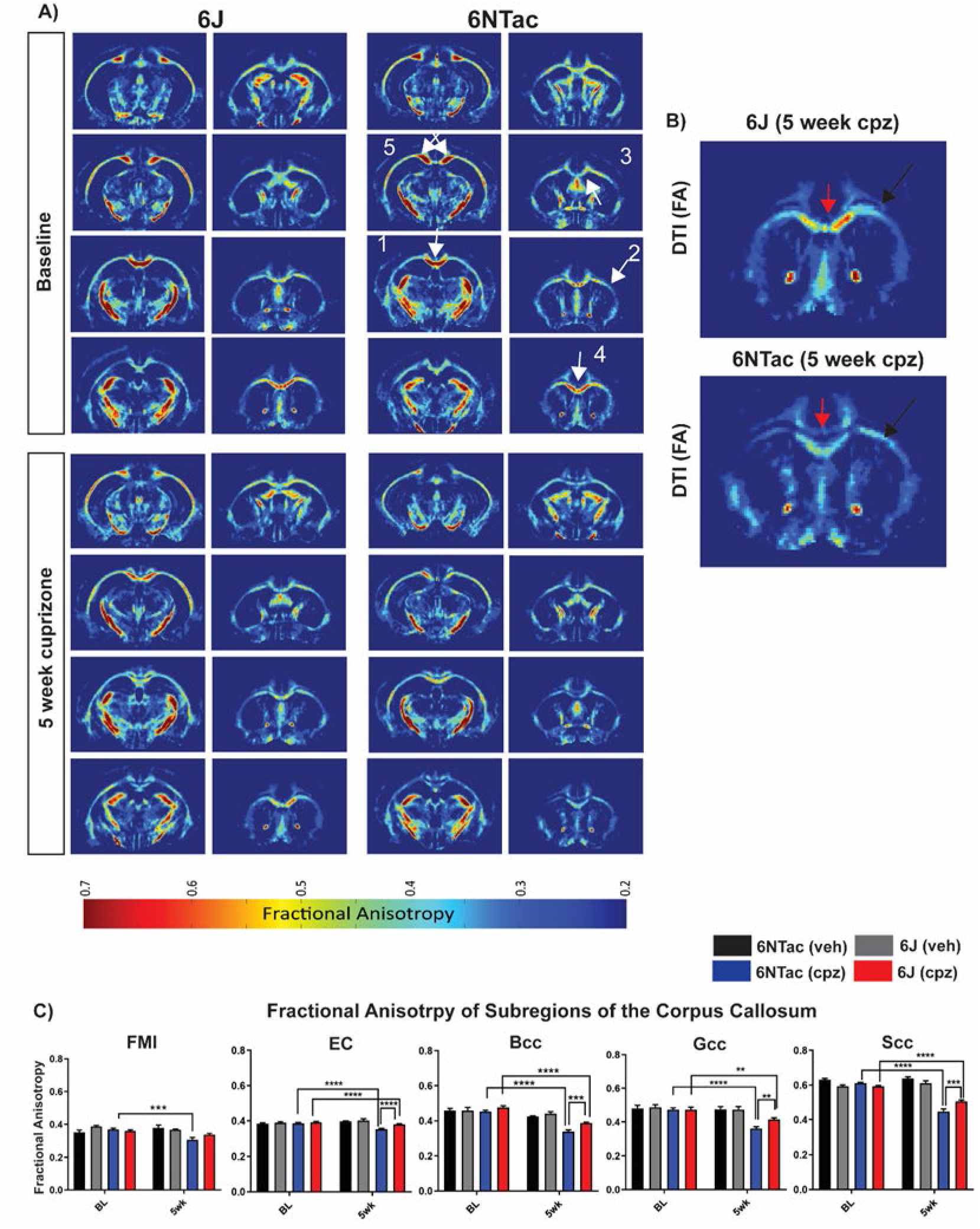
A) Representative images of DT-MRI fractional anisotropy analysis of C57BL/6NTac and C57BL/6J mice at baseline and 5-week cpz treatment. Color scale indicates the FA intensity (low=0.2 to high=0.7). B) Images of FA 6J (5 week cuprizone) and 6NTac (5 week cuprizone) C) Fractional anisotropy (FA) measurements of subregions of the Corpus Callosum (CC) including the *forceps minor, external capsule, body, genu and the splenium,* imaged at baseline (before dosing) and at 5 weeks dosing of either vehicle (veh) or cpz (cpz), respectively. White arrows indicate regions of the cc 1) Body of CC, 2) external capsule, 3) forceps minor, 4) genu and 5) the splenium. Red arrows indicate the body of the CC and black arrows indicate the external capsule. BL is baseline, before any treatment occurred. Data represented mean ±SEM. Analysis carried out by Two-way ANOVA with Tukey’s multiple comparisons post hoc (*, p<0.05; **, p<0.01; ***, p<0.001). Animal groups: n= 12 6J+cpz, n= 12 6NTac+cpz, n=4 6J+vehicle, n=4 6NTac+vehicle.

### 2.3 Demyelination and cellularity differences between C57BL/6J and C57BL/6NTac strains

Sections were semi-quantitatively evaluated for decrease in LFB staining (indicative of demyelination) and for increased cellularity in the corpus callosum (indicative of reactive gliosis) *(Supplementary Table 2).* Cpz treatment induced marked to severe demyelination at different levels of the CC as characterized by decreased to complete absence of LFB staining in both C57BL/6Ntac and C57BL/6J groups (Figure 3B-D). Overall, no major difference in severity of demyelination was noted between substrains of mice tested by luxol fast blue stain and pathology scoring. It should be noted that the severity of demyelination varies between different regions of the CC, suggesting potential sub-regional sensitivities of myelinating cells to cpz treatment. The severity of demyelination was associated with a concomitant minimal to moderate increase in cellularity in the CC in C57BL/6NTac in comparison C57BL/6J mice after 5 weeks of CPZ treatment (Figure 3D). The increase in cellularity comprised primarily of microglial cells in response to the demyelination as confirmed by Iba1 IHC (Figure 3D). The changes in cellularity observed were moderate, although they may indicate a reduction in gliosis response in these mice. One animal recorded no response to cpz, with normal myelination and no cellular response and was thus excluded from further downstream analysis.

**Figure 3:**
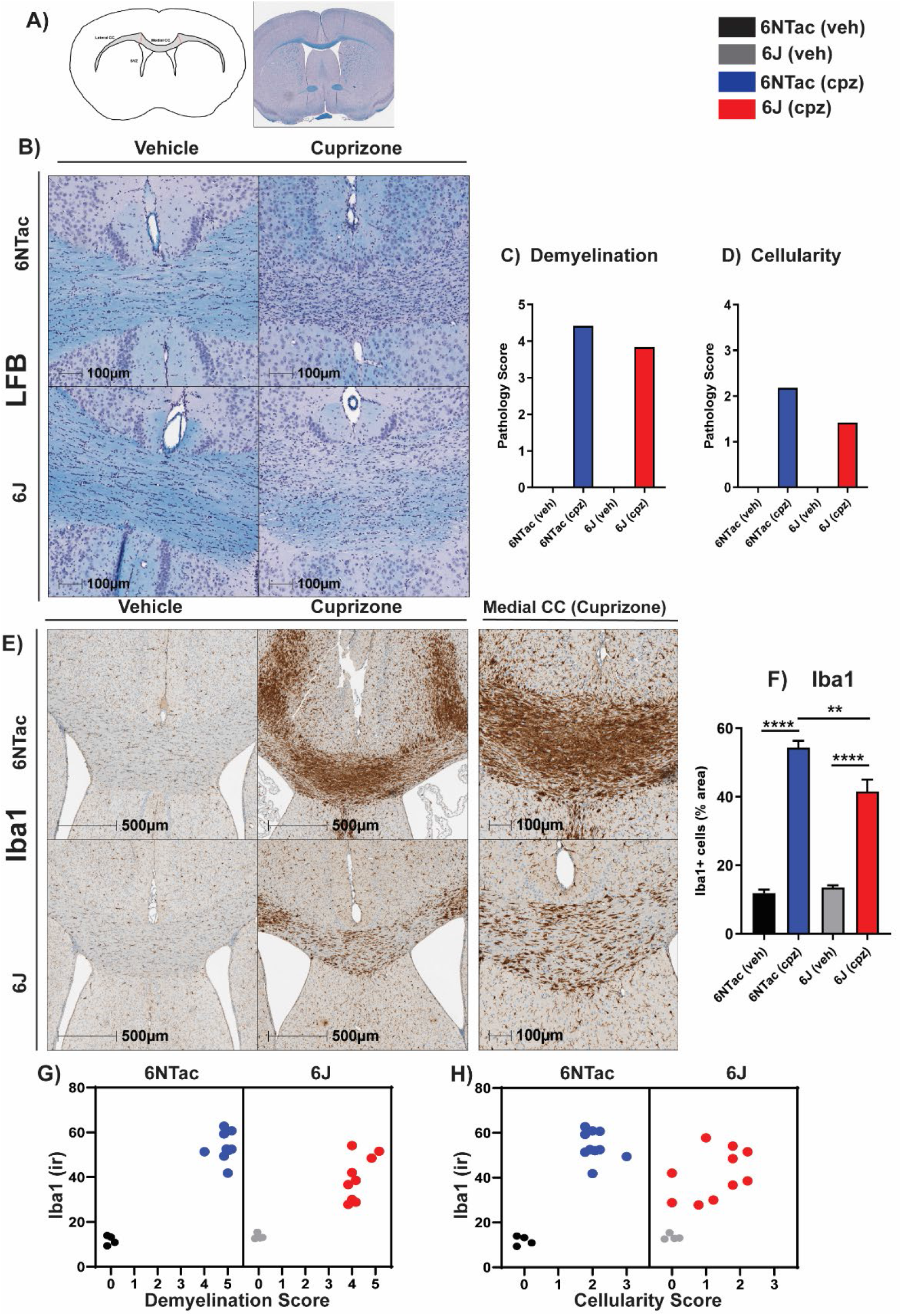
A) Representative images of LFB stained sections showing the genu of the Corpus Callosum (CC). Blue stain depicts myelinated fibers and purple nuclear counter staining. B, C) Qualitative histopathology scoring for demyelination and cellularity. LFB demyelination was scored according to a pathology severity grade from score 1-5, where 1 = minimal and 5 = almost complete loss of LFB staining. Cellularity was scored in a similar way, with 1 = minimal and 5 = severe increase in cellularity. In each case WNL (0) denoted within the normal limits i.e. no change. Bar graphs show group mean score, vehicle (veh) n=4, cpz (cpz) n=12 mice per group. D) Iba1 representative images in the genu of the CC; *F (3, 24) =42.75;* p<0.0001. E) Quantification of Iba1+ cells in the CC. Animal groups; n=11 6J+cpz, n=12 6NTac+cpz and n=4 6J+vehicle, 6NTac+vehicle per group. Data represented group mean ±SEM. F, G) Scatter plots of Iba1 immunoreactivity (ir) in comparison to demyelination and cellularity score as scored on LFB sections.

### 2.4 Microglial response after cpz treatment

Microglia/macrophage response as measured by Iba1 staining and was dramatically increased in response to 5-weeks cpz treatment in the CC of treated animals vs. vehicle groups independent of substrain (Figure 3E, F). Quantification of Iba1 area revealed a significant difference in immunoreactivity between the cpz groups, C57BL/6NTac (54% area covered) and C57BL/6J (41% area covered) mice *(F (3, 24) = 42.75, p<0.0001* Figure 3E). This difference suggests that C57BL/6J mice may have a diminished microglial/macrophage response to cpz toxin. There was no change in Iba1 immunoreactivity in vehicle treated groups between the two substrains. A comparison of Iba1 immunoreactivity versus pathology scores was plotted for the two strains (Figure 3G, H). The scatter plots show that high demyelination scores correlate with high Iba1 immunoreactivity (Figure 3F) and this is also true for the cellularity score (Figure 3G). Interestingly, C57BL/6NTac mice exhibited more consistent cellularity scores and Iba1 response than C57BL/6J mice in CC lesions at peak of demyelination, which may illustrate that C57BL/6NTac mice respond to the toxin more robustly at the cellular level.

### 2.5 Cellular functional responses between C57BL/6NTac and C57BL/6J mice after cpz

To examine the extent of myelin debris generation and as an indirect evaluation of microglia phagocytic function, we used an antibody specific for degraded myelin basic proteins (dMBP). Data showed that C57BL/6NTac mice had significantly higher levels of dMBP (puncta intensity) in the CC in response to cuprizone treatment relative to vehicle animals (*F (3, 22) = 14.4, p<0.0001,* Figure 4A, B). C57BL/6J did not display significant differences in dMBP puncta with cpz treatment relative to vehicle treated groups (Figure 4B). Vehicle-treated C57BL/6NTac and C57BL/6J strains showed similarly low levels of dMBP while cpz treated C57BL/6NTac mice had significantly higher absolute levels of dMBP immunoreactivity compared to the cpz-treated C57BL/6J mice (Figure 4A, B). The proliferation marker Ki-67 was used to examine changes in CC lesion cell numbers in response to cpz treatment (Figure 4C). C57BL/6NTac mice exhibit a significantly higher proliferation response to cpz induced damage versus C57BL/6J mice (*F (3, 23) = 20.03, p<0.0001,* Figure 4D). It should be noted that, proliferating cells may represent microglia, astrocytes and/or oligodendrocyte precursors.

**Figure 4:**
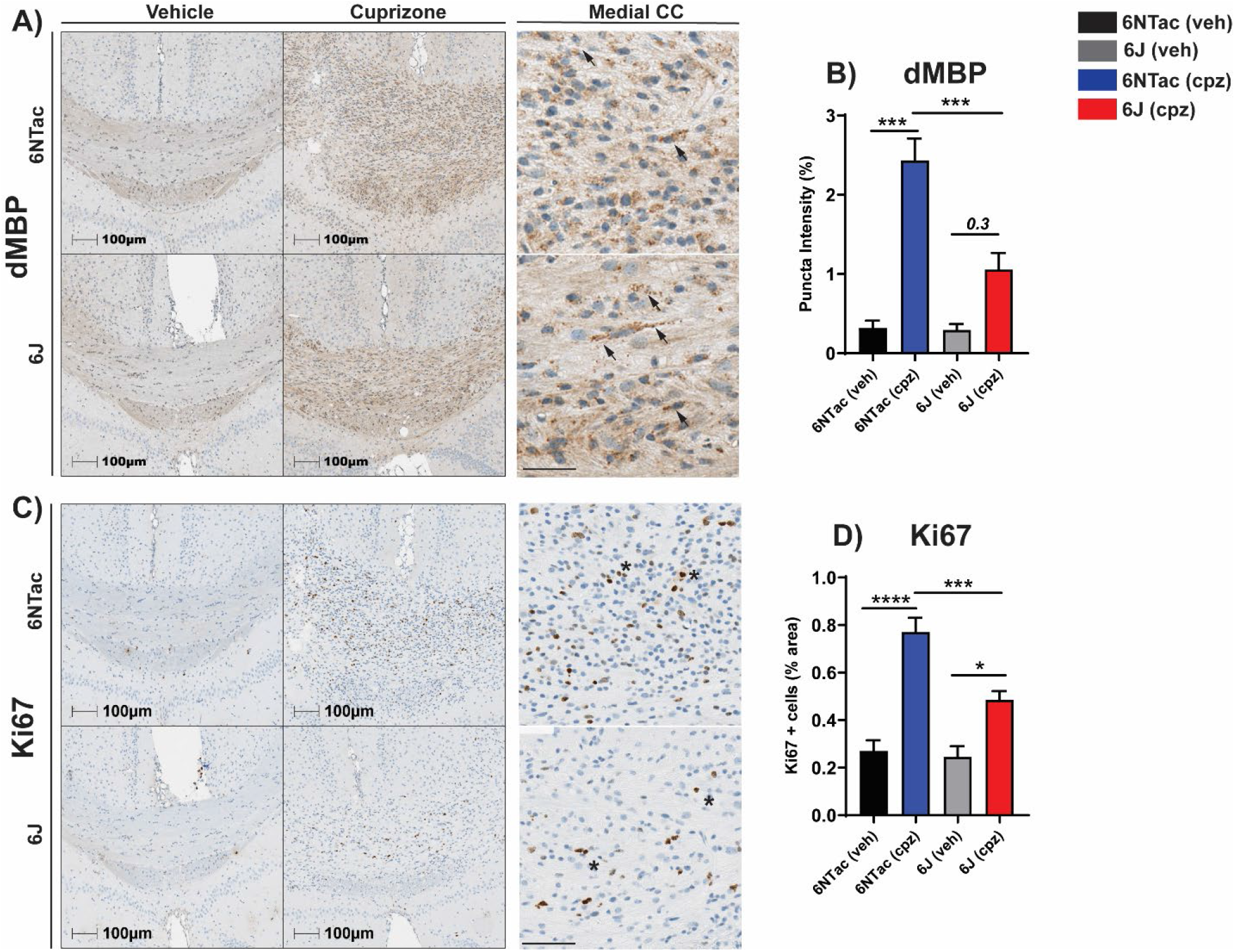
A) Degraded myelin basic protein immunostaining in the Corpus Callosum (CC) of vehicle (veh) and cpz (cup) C57BL/6NTac and C57BL/6J mice. High magnification inset of the medial CC showing dMBP+ puncta after cpz treatment. C) Quantification of dMBP puncta in the CC as % intensity; *F (3, 22) = 14.4; p<0.0001.* B) Ki-67 representative images with high magnification inset of the medial CC showing Ki-67+ cells after cpz treatment. D) Quantification of Ki-67+ cells in the CC as % area. Data represented mean ±SEM. Analysis carried out by one-way ANOVA with Tukey’s multiple comparisons post hoc (*, p<0.05; **, p<0.01; ***, p<0.001). Animal groups; n=11 6J+cpz, n=12 6NTac+cpz and n=4 6J+vehicle, 6NTac+vehicle per group.

### 2.6 Oligodendrocyte cell population is unchanged between strains after cpz treatment

Olig2 is a pan oligodendrocyte cell linage marker that identifies oligodendrocyte progenitor cells, premyelinating cells and fully mature oligodendrocytes. Representative images show the expected disruption of oligodendrocyte cytoarchitecture in both mouse strains after cpz treatment in comparison to vehicle controls *(Supplementary Figure 2*). In vehicle treated animals, Olig2+ cells are uniformly arrayed along fibers in the CC but after cpz treatment, the cells appear randomly distributed throughout the CC (*Supplementary Figure 2*). Quantification of Olig2+ cells in the CC showed that there were no significant differences in Olig2+ cell numbers observed between substrains after cpz treatment *((F (3, 23) = 0.2273, p=0.8765, Supplementary Figure 2*).

## 3. Discussion

Our comparative analysis demonstrated that C57BL/6NTac and C57BL/6J substrains, presumed to be highly similar based on their common origins and their indistinguishable baseline characteristics, display differential sensitivity to cpz toxin at the level of body weight, DTI changes and multiple readouts of myelin histopathology and microglia response. The C57BL/6NTac mice were found to be more susceptible to the toxin challenge over a 5-week period. At the CNS structural level, DTI is a powerful translational neuroimaging technique, which allows longitudinal intravital image acquisition and sensitive detection of changes of brain structures, myelin integrity, axonal injury and cellular infiltration. DTI measurements, including fractional anisotropy and mean diffusion. Fractional anisotropy is a diffusion tensor property, which is directionally dependent and sensitive to structural coherence. Mean diffusion, on the other hand, is orientation independent and would report more generalized changes such as hindrance of the overall diffusion (e.g. reduction due to the myelin debris accumulation) or increase due to increased diffusion space. While the such changes can affect both parameters, FA is normally more sensitive in the context of demyelination. Cpz treatment largely affects the white matter structures of the CNS with overt changes observed in the CC and surrounding subregions. It should be noted that other brain structures can also be susceptible to challenge, for example, the hippocampus, cortex and cerebellum, which were not addressed in this present study (Koutsoudaki et al., 2009, Sachs et al., 2014, Sen et al., 2020). The above-mentioned studies used different doses, routes of administration, exposure times and mouse strains, which are all important factors to recognize as each experimental paradigm will yield very different results. In the present study, DTI illustrated that the C57BL/6NTac mice had significant effects after cpz treatment with significant decreases in fractional anisotropy read-outs in major subregions of the CC. At the histology level, both strains exhibited comparable demyelination as indicated by pathology evaluation on LFB stained brain sections, while the cellularity scores were moderately different, which may indicate that the C57BL/6NTac mice exhibit different cellular responses than C57BL/6J after toxin challenge. Further analysis of Iba1+ microglia indicated a robust response in CC regions but with significant increases in immunoreactivity in C57BL/6NTac versus C57BL/6J mice, thus demonstrating a varied central innate immune response in the C57BL/6NTac strain. That may result from multiple variable such as increased level of myelin damage, higher sensitivity of microglia to toxin induced damage or simply greater resilience of 6J mice. This would be important when studying microglial targets, for example, TREM2, or cellular responses to the disease niche as the results could be confounded by inappropriate strain selection (Ohrfelt et al., 2016, Gratuze et al., 2018). The data would suggest that the C57BL/6NTac background strain is more suitable to monitor cpz mediated demyelination. Degraded myelin basic protein could be interpreted as an indirect read-out of clearance of myelin debris. The cellularity scores, Iba1 and dMBP immunoreactivity were compared for each strain and treatment group. The levels of dMBP immunoreactivity were significantly different but this covers only 1-2.5% of the CC after 5 weeks dosing, and it appears that both strains’ cellular response can clear the debris. Oligodendrocytes are vulnerable to CPZ and this study showed that the number of Olig2+ cells does not differ after 5-week treatment; however, the structural architecture of the CC myelinating fiber is dramatically different between vehicle- and cpz-treated animals. It should be noted that Olig2 a pan marker oligodendrocyte and does not discriminate between all oligodendrocyte states e.g. OPCs. However, the Olig2 antibody has been routinely used in studies to examine oligodendrocytes in *in vivo* studies, particularly in the cuprizone toxin model (Islam et al., 2009, Ye et al., 2013). In addition, while we did not address gender in this study, male mice were used in cuprizone treatment studies. Female mice may be more resistant to cuprizone-toxin induced demyelination (Hibbits et al., 2009, Steelman et al., 2012). However, one recent publication suggests that female mice may be sensitive, however, in our studies we used male mice (Almuslehi et al., 2020).

Simon et al. (2013), carried out a comprehensive genomic and phenotypic analysis between C57BL/6J and C57BL/6NTac untreated healthy mice to examine any difference that may underlie genetic mechanisms (Simon et al., 2013). This study showed that there are sets of candidate genes that are significantly different between the substrains. The group also noted differences in the strains from separate research facilities and documented that in naïve states, these strains are very diverse. Across the four centers involved in the study, C57BL/6NTac and C57BL/6J mice differed in several areas including behavioral (startle response, locomotor activity, grip strength), physiological (cardiovascular characteristics, metabolic parameters) and in clinical chemistry parameters (Simon et al., 2013). Some of the underlying spontaneous mutations responsible for the heritable phenotypic differences between C57BL/6NTac and C57BL/6J substrains, which have arisen due to genetic drift, previously have been identified (Radulovic et al., 1998, Stiedl et al., 1999, Roth et al., 2002, Mekada et al., 2009, Mattapallil et al., 2012, Fernando et al., 2016, Fontaine and Davis, 2016). The Jax website also discusses the topic that there is no such thing as a B6 mouse and the present study illustrates this further (https://www.jax.org/news-and-insights/jax-blog/2016/june/there-is-no-such-thing-as-a-b6-mouse). Previous studies have reported on strain differences in cpz models between CD1 and C57BL/6 mice, noting significant differences in demyelination, behavior and cell responses (Yu et al., 2017). In 2009, Taylor et al investigated the SJL strain in the cpz paradigm and found this to be less susceptible to demyelination in the CC compared to the established C57BL/6 strain (Taylor et al., 2009). Additional strain comparison papers include BALB/cJ versus C57BL/6 mice, where cortical demyelination in BALB/cJ mice treated with 0.2% cpz for 6 weeks is incomplete in comparison to the C57BL/6 strain (Skripuletz et al., 2008). In some published articles describing strain differences to the effects of cuprizone, the methods sections often do not indicate the substrain used e.g. stating only C57BL/6 mice. To date, there have been no comparative data reported between C57BL/6J and C57BL/6NTac mice in the cpz model and this study is the first to elucidate strain-based differences. Consistent with the increased stroke vulnerability reported by Nowak & Mulligan (2019), the present study shows that C57BL/6NTac mice are more susceptible to cpz toxicity in comparison to C57BL/6J mice which is an important factor to consider when choosing the correct strain background for transgenic lines. The use of transgenic mice in investigating loss-or gain-of-function models in the context of myelination and axonal pathology is highly informative (Nowak and Mulligan, 2019). Substrain genetic background might influence and/or modulate oligodendrocyte maturation/stability and dynamics of myelination upon cpz sensitivity and the pattern of inducible demyelination, which is important to consider when evaluating a potential disease modifying target or drug candidate.

## 4. Conclusion

The C57BL/6 mouse model strain is a powerful research tool that has been invaluable to the progress of biomedical research, especially to neuroscience research. Researchers are using the B6 mouse to gain deeper insights into numerous disorders and diseases. It is important to know the relative sensitivity between strains when developing an experimental model and choosing the appropriate strain background for the genetically engineered strain. In addition, choosing the appropriate mouse strain to study a therapeutic intervention window is vital in producing reliable, meaningful data and preventing data from being misinterpreted. Thus, it is important to comprehend the window of biology in which your candidate molecule may modulate physiology, resulting in beneficial or harmful effects in your model strain. To conclude, careful consideration should be taken when designing studies using the cpz model but also for additional challenge-based models.

## 5. Experimental procedure

### 5.1 Animals

Male C57BL/6NTac mice (Taconic Laboratories, Germantown, NY), and C57BL/6J mice (Charles River Laboratory, Germany) at 10-12 weeks of age were treated with either vehicle (0.5% methylcellulose in water) or 100 mg/kg of cpz formulated in vehicle twice daily (b.i.d) by oral gavage for 5 weeks (200 mg/kg total dose/per day). Animal groups included *C57BL/6J (veh) n=4; C57BL/6NTac (veh) n=4; C57BL/6J (cpz) n=12; C57BL/6NTac (cpz) n=12.* The in-life components and termination procedures for this study were conducted by Charles River Laboratories (Kuopio, Finland).

### 5.2 Cuprizone treatment

Cuprizone (Fluka/Sigma-Aldrich Ca no. 14690) dosing suspension for the oral gavages was formulated in 0.5% methylcellulose in water (400 cp; Sigma-Aldrich Cat: M0262). All mice received a total cpz dose of 200 mg/kg/day, (100 mg/kg per dosing, every 10-12 hours) or the corresponding volume of vehicle. In the present study, cpz was administered via oral gavage using a dosing protocol that standardized daily administration to each animal based on the individual bodyweight. Oral gavage allows the investigator to control for administration and timing of delivery, is more reproducible than chow options. Thus, animal numbers can be reduced while still sufficiently/appropriately powering studies. An internal dose response cpz study was conducted (data not shown) and showed robust effects on body weight loss, demyelination and gliosis at lower doses (200 mg/kg/day) compared to previously published data (Zhen et al., 2017).

### 5.3 Body weight measurements

Body weight (BW) loss is a phenotypic outcome with cpz administration in mice. Body weights were measured daily commencing on the day of first dosing (Day 1). The measurements took place each day prior to morning dosing of cpz to allow for adjustment of cpz dose. Weight loss is a clinical parameter of this model; therefore, nutrient gel was supplied to animals after 15% BW loss was observed. Mice that reached or exceeded 25% BW loss were removed from the study.

### 5.4 Diffusion Tensor Magnetic Resonance Imaging (DT-MRI)

MRI acquisitions were performed for all available mice at baseline and at 5 weeks of cpz/vehicle dosing (demyelination phase). For DT-MRI, a horizontal 11.7T magnet with a bore size of 160 mm was used. The magnet was equipped with a gradient set capable of maximum gradient strength of 750 millitesla per meter (mT/m) and was interfaced to a Bruker Avance III console. A volume coil was used for transmission and a surface phased array coil for receiving. Mice were anesthetized using isoflurane, fixed to a head holder and positioned in the magnet bore in a standard orientation relative to gradient coils. After acquisition of fast localizer images, DT-MRI was performed using 4-segment EPI sequence with 30 diffusion directions (TE/TR =23.5/4000 ms, b-values 0 and 970 s/mm2). Field-of-view of 12.80 x 10.24 mm^2^ (with saturation slice) was used with matrix of 160 x 128, resulting in 80 microns in-plane resolution. Fifteen 0.6 mm slices were acquired with 6 averages. Preprocessing of DT-MRI-data consisted of eddy-current correction and brain masking. Diffusion tensor as a measure of white matter integrity was calculated using FSL and resulting fractional anisotropy and mean diffusion maps were processed with manual ROI-analysis for the following anatomical structures; *forceps minor of the corpus callosum (fmi), genu of corpus callosum (gcc); body of corpus callosum (bcc); splenium of corpus callosum (scc); external capsule (ec); anterior commissure anterior part (aca); internal capsule (ic); optic tract (opt); cerebral peduncle (cp).*

### 5.5 Histology

#### 5.5.1 Tissue preparation and Luxol Fast Blue staining for evaluation of myelin integrity

At the end of the study, all mice were terminally euthanized with an overdose of CO_2_ and transcardially perfused with phosphate buffered saline (PBS) followed by 10% neutral buffered formalin. Brains were immersion fixed for 24-48h in 10% formalin. Tissue was trimmed into 2-mm tissue slabs, paraffin-embedded and serially sectioned in the coronal plane on the microtome. Brain sections cut at 12μm were, deparaffinized, hydrated and stained by placing slides in LFB (0.1%) solution for 30 min at 60°C (Margolis and Pickett, 1956). Semi-quantitative histomorphology assessment was performed by a certified veterinary pathologist on LFB and hematoxylin counter-stained sections of the sections from all animals. Using the concurrent control group animals as baseline, demyelination and increase in cellularity were scored using a five point severity grading scheme, ranging from minimal (Grade 1, ~ <10% change from normal) to severe (Grade 5; ~ >80% change from normal) with ranges of 10 to 20% change (Grade 2, mild), 20 to 40% change (Grade 3, moderate) and 40 to 80% change (Grade 4, marked) in between. In each case WNL (0) denoted within the normal limits (i.e., no change).

#### 5.5.2 Immunohistochemistry

For immunohistochemistry, the same brains for LFB were sectioned at 5 μm serial sections. All tissue samples were processed on the Ventana BenchMark ULTRA system (Roche Tissue Diagnostics). Information on antibodies used are in Table 1. Briefly, sections were de-waxed and rehydrated. Depending on the antibody, different antigen retrieval methods were incorporated. Sections were stained with selected antibody (Table1) in 10% normal donkey serum, 0.3% sodium azide and 0.1% Triton X-100 solution. Antibody detection was carried out using the appropriate secondary antibody and counterstained with hematoxylin and eosin stain and Nissl stain. Slides were then dehydrated and cover-slipped.

**Table 1:** Antibodies used for immunohistochemistry on paraffin embedded sections.

| Antibody | Company | Antigen Retrieval | Species |
| --- | --- | --- | --- |
| Iba-1<br>(microglia/macrophage marker) | Wako<br>019-19741 | Tris-EDTA buffer pH 7.8 | Rabbit;<br>1 in 2000 dilution |
| dMBP<br>(myelin basic protein) | EMD Millipore<br>ab5864 | Tris-EDTA buffer pH 7.8 | Mouse;<br>1 in 4000 dilution |
| Ki-67<br>(proliferation marker) | Abcam<br>ab15580 | Tris-EDTA buffer pH 7.8 | Rabbit;<br>1 in 1000 dilution |
| Olig2<br>(oligodendrocyte marker) | Abcam<br>ab109186 | Citrate-based buffer pH 6.0 | Rabbit;<br>1 in 1000 dilution |

### 5.6 Quantification and statistical analysis

Whole slide images were captured on an Aperio Digital Pathology XT Slide Scanner (Leica Biosystems) at 40x magnification. Quantification was carried out using Halo™ image analysis platform (Indica Labs) and algorithm “Area Quantification”. Two sections containing the corpus callosum regions (CC) per animal, per group were analyzed. The CC was outlined manually using reference from mouse brain atlas (bregma 0.75mm) and measurements/analyses (% marker positive area) were carried out on this region of interest. This included lateral and medial portions of the CC as outlined in histology schematic (Figure 3A). Statistical analyses were carried out using GraphPad Prism software version 8.1.1. Data were represented as mean values ± the standard error of the mean (SEM). Differences were determined between C57BL/6J and C57BL/6NTac mice using either unpaired student’s t-test or one-way or two-way ANOVA analysis followed by multiple-comparison post hoc test where appropriate. Differences were significant only for *\*\*\*p < 0.001, **p < 0.01,* and *\*p < 0.05.*

## Acknowledgements

The authors would like to thank Christopher Ware and Gain Robinson for histology recommendations and consulting on the DT-MRI analysis, respectively.

## Authors’ contributions

EOL contributed to study conception and design, preformed the experiments, collected data, performed data analysis, and wrote the manuscript. A-MK, KL, and JK conducted the in-life portion of the study. SB contributed to study conception and design, method development, and manuscript revisions. CM contributed to study conception and design and manuscript revisions. MK lead study approval and manuscript revisions. GZ carried out pilot histology experiments. BJ, PM, TT AP, NTG Coordination of pathological endpoints and FFPE tissue processing and manuscript revisions. TR and GV directed studies at CRL Finland. All authors read and approved the final manuscript.

## Ethic approval

Principles of laboratory animal care were followed, and all studies were previously approved by the Institutional Animal Care and Use Committee and were performed in accordance to the Guide for the Care and Use of Laboratory Animals as adopted and promulgated by the National Institutes of Health (Library of Congress Control Number 2010940400, revised 2011).

## Availability of data and materials

The datasets used and/or analyzed during the current study are available from the corresponding author upon acceptance.

## Consent for Publication

Not applicable

---

**Supplementary Figure 1**: A) Mean diffusion analysis of DT-MRI including subregions of the Corpus Callosum (CC); Body of CC, Forceps Minor of CC; External Capsule; Genu of CC and splenium of CC. Data represented mean ±SEM. Analysis carried out by one-way ANOVA with Tukey’s multiple comparisons post hoc (*, p<0.05; **, p<0.01; ***, p<0.001). For both strains 6NTac and 6J animal groups are n= 11 cpz and n=4 vehicle per group.

**Supplementary Figure 2**: A) Panel of images of oligodendrocyte (Olig2+ cells) in the CC in vehicle and cpz treated mice. Representative images of Olig2 with high magnification inset of the medial CC showing cells after cpz treatment. B) Quantification of Olig2+ cells in the CC as percentage area (*F (3, 23) = 0.2273, p=0.8765).* Data represented mean ±SEM. Analysis carried out by one-way ANOVA with Tukey’s multiple comparisons post hoc. For both strains 6NTac and 6J animal groups are n= 11 cpz and n=4 vehicle per group.

**Supplementary Table 2:** Summary of histomorphological changes in LFB stained brain sections. Severity grade 1-5 was used (1 = minimal; 5 = almost complete loss of staining) b. Severity grade 1-5 was used (1 = minimal; 5 = severe increase in cellularity) WNL = within normal limit.

**Table 2.**

| **Genotype** | **Treatment** | **Decreased LFB staining<sup>a</sup>** | **Increased cellularity<sup>a</sup>** |
| --- | --- | --- | --- |
| C57BL/6NTac | CPZ | 5 | 2 |
| C57BL/6NTac | CPZ | 4 | 2 |
| C57BL/6NTac | CPZ | 5 | 2 |
| C57BL/6NTac | CPZ | 5 | 2 |
| C57BL/6NTac | CPZ | 5 | 2 |
| C57BL/6NTac | CPZ | 4 | 2 |
| C57BL/6NTac | CPZ | WNL | WNL |
| C57BL/6NTac | CPZ | 5 | 2 |
| C57BL/6NTac | CPZ | 5 | 3 |
| C57BL/6NTac | CPZ | 5 | 3 |
| C57BL/6NTac | CPZ | 5 | 2 |
| C57BL/6NTac | Vehicle | WNL | WNL |
| C57BL/6NTac | Vehicle | WNL | WNL |
| C57BL/6NTac | Vehicle | WNL | WNL |
| C57BL/6NTac | Vehicle | WNL | WNL |
| C57BL/6J | CPZ | 4 | 1 |
| C57BL/6J | CPZ | 4 | 1 |
| C57BL/6J | CPZ | 5 | 2 |
| C57BL/6J | CPZ | 5 | 2 |
| C57BL/6J | CPZ | 4 | 2 |
| C57BL/6J | CPZ | 4 | 1 |
| C57BL/6J | CPZ | 4 | WNL |
| C57BL/6J | CPZ | 4 | 2 |
| C57BL/6J | CPZ | 5 | 2 |
| C57BL/6J | CPZ | 4 | 2 |
| C57BL/6J | CPZ | 4 | WNL |
| C57BL/6J | CPZ | 3 | 2 |
| C57BL/6J | Vehicle | WNL | WNL |
| C57BL/6J | Vehicle | WNL | WNL |
| C57BL/6J | Vehicle | WNL | WNL |
| C57BL/6J | Vehicle | WNL | WNL |
| a. A five point severity grading scheme, ranging from minimal (Grade 1, ~ <10% change from normal) to severe (Grade 5; ~ >80% change from normal) and with ranges of 10 to 20% change (Grade 2, mild), 20 to 40% change (Grade 3, moderate) and 40 to 80% change (Grade 4, marked) in between |  |  |  |

