## Supplementary figures and images for "C57BL6 mouse substrains demonstrate differences in susceptibility to the demyelinating effects of Cuprizone toxin"

### Supplemental Figures 1&2

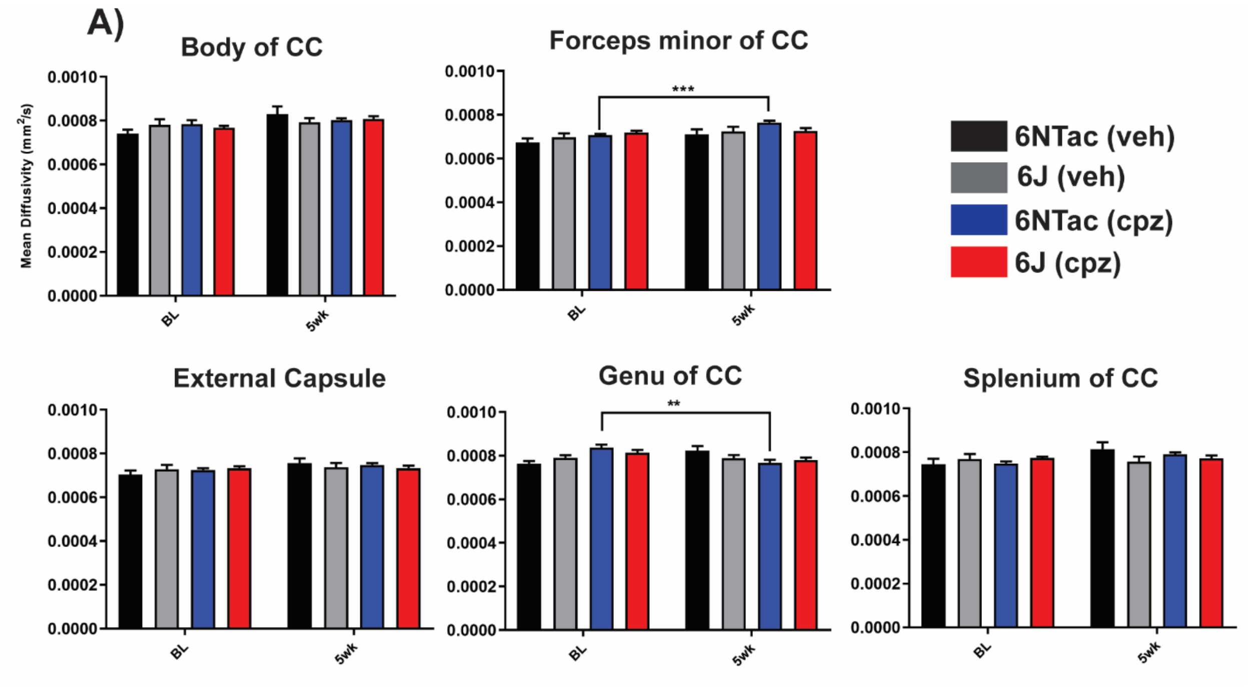


Supplementary figure 1

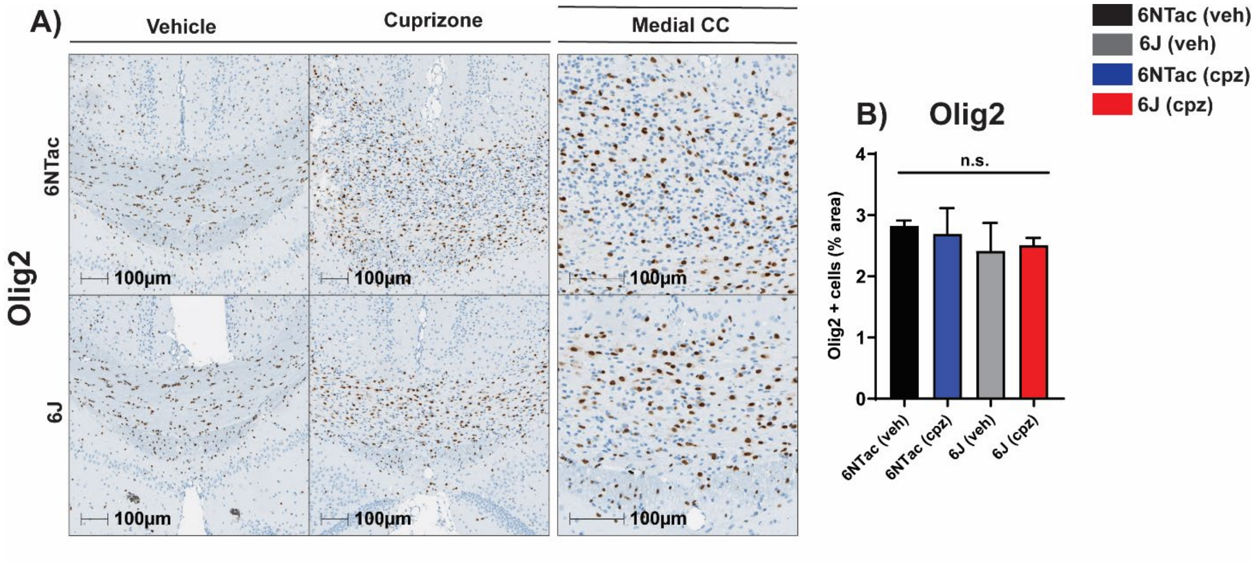


Supplementary figure 2
